# *In vivo* structures of the *Helicobacter pylori cag* type IV secretion system

**DOI:** 10.1101/195685

**Authors:** Yi-Wei Chang, Carrie L. Shaffer, Lee A. Rettberg, Debnath Ghosal, Grant J. Jensen

## Abstract

The bacterial type IV secretion system (T4SS) is a versatile nanomachine that translocates diverse effector molecules between microbes and into eukaryotic cells. Using electron cryotomography, here we reveal the molecular architecture of the cancer-associated *Helicobacter pylori cag* T4SS. Although most components are unique to *H. pylori*, the *cag* T4SS exhibits remarkable architectural similarity to previously studied T4SSs. When *H. pylori* encounters host cells, however, the bacterium elaborates rigid, membranous tubes perforated by lateral ports. Dense, pilus-like rod structures extending from the inner membrane were also observed. We propose that the membrane tubes assemble out of the T4SS and are the delivery system for *cag* T4SS cargo. These studies reveal the architecture of a dynamic molecular machine that evolved to function in the human gastric niche.

## Introduction

The type IV secretion system (T4SS) is a remarkably versatile molecular machine present in nearly all bacterial phyla and some archaeal species (*1*). Bacteria utilize T4SS to interact with prokaryotic and eukaryotic cells and to export an incredibly diverse repertoire of substrates (*2*). In most cases, T4SS activity is contact-dependent and results in delivery of nucleoprotein complexes and protein effectors directly into the target cell cytoplasm. By facilitating the exchange of genes and proteins among microbial populations and across kingdoms of life, the T4SS has accelerated bacterial evolution and resulted in species that thrive in diverse environments, including within plant and animal hosts (*1, 3, 4*).

T4SSs are used by a variety of pathogens during host colonization, including the gastric bacterium *Helicobacter pylori*. *H. pylori* may harbor up to four T4SSs, including the *cag* pathogenicity island (*cag*PAI)-encoded T4SS (*cag* T4SS) (*5, 6*); the *comB* T4SS that mediates DNA uptake from the extracellular environment (*7*); and two less well-characterized T4SSs, *tfs3* and *tfs4*, which are hypothesized to function in horizontal DNA transfer between bacteria (*8*). *H. pylori* exploit the *cag* T4SS to translocate a variety of effector molecules into gastric epithelial cells, including the oncoprotein CagA, fragments of peptidoglycan, chromosomally-derived DNA, and the lipopolysaccharide (LPS) biosynthesis metabolite heptose-1,7-bisphosphate (*5, 9-11*). These translocated effector molecules activate components of the innate immune system and dysregulate signaling pathways that significantly augment the risk of gastric cancer (*12, 13*).

Elegant studies analyzing the prototypical *vir* T4SS harbored by the phytopathogen *Agrobacterium tumefaciens* have provided valuable insight into T4SS biogenesis and function (*1, 14*). In addition, recent work has provided architectural and structural information about several different T4SSs including the *Escherichia coli* conjugation *tra* T4SS (*15, 16*), the *A. tumefaciens vir* T4SS (*17*), the *Legionella pneumophila dot/icm* T4SS (*18, 19*), and the *H. pylori cag* T4SS (*20*). Among these, the *dot/icm* and *cag* effector translocator systems have many more genes than the *tra* and *vir* DNA-translocating systems, including many without obvious homologs in other bacteria (*18-20*). A so-called “core complex” of the *cag* T4SS has been purified and consists of Cag3, CagT, CagM, and two constituents that are orthologous to the VirB9/TraO (CagX) and VirB10/TraF (CagY) subunits of other systems (*6, 20*). When *H. pylori* contact the gastric cell surface, the bacterium produces filamentous structures that are dependent on multiple *cag* genes and have been termed *cag* T4SS pili (*21-26*). While *H. pylori* strains that fail to produce *cag* T4SS pili are unable to translocate cargo to host cells (*22, 23, 25*), the exact role of these filaments in *cag* T4SS activity and the relationship between the *cag* T4SS and the filaments remain unclear. In the current study, we applied electron cryotomography (ECT) to image frozen-hydrated *H. pylori* co-cultured with human gastric epithelial cells. We report structures of the intact *cag* T4SS *in vivo* and describe membranous tubes elaborated by *H. pylori* in response to the host cell. Together, these results suggest new hypotheses about the mechanism of the *cag* T4SS and roles of its components.

## Results

### *H. pylori* develop membranous tube-like appendages in response to the host cell

Since T4SS activity is stimulated by direct host cell contact, we sought to visualize the *cag* T4SS by imaging *H. pylori* in co-culture with human gastric epithelial cells. In order to avoid interference from flagella or flagellar motors in the analysis, we selected the *cag*PAI-positive, non-flagellated *H. pylori* strain 26695 for our studies. We cultured gastric epithelial cells on electron microscopy grids, infected the grid-adherent monolayers with *H. pylori,* and plunge-froze the co-culture sample to preserve cellular features in a near-native state. We recorded ECT tilt-series (*27*) of regions of the sample where the bacteria were in direct contact or close proximity to epithelial cell elongations (Fig. 1A, B). In approximately 5% of the tomograms, we observed striking membranous tubes extending from the outer membrane of *H. pylori* cells (Fig. 1C, D) which were not observed in the absence of gastric epithelial cells (n = 464, Fig. S1). While this indicates that the tubes assemble in response to the host cell, we did not observe direct interaction of individual tubes with gastric epithelial cell surfaces (though we cannot rule out the possibility that longer tubes touched host cell surfaces beyond the imaging area of individual tomograms). Putative LPS densities, as observed on the cell surface, were clearly visible on the periphery of tube cross-sections, as were both leaflets of the membrane bilayer (Fig. 1D, inset). A thin layer of periodic densities lined the interior of the tubes, intimately associated with the inner leaflet of the bilayer, suggesting the presence of a regular protein support scaffold (Fig. 1E). The tubes appeared rigid and had membrane-outer-surface (not including LPS) and inner-channel (inside surface of scaffold) diameters of 37- and 22-nm, respectively (Fig. S2). Many tubes displayed pipe-like ports (median dimeter 10 nm) along their lengths (Fig. 1F-I). In some cases, ports appeared to induce small bends in the tube (Fig. 1J), as if by wedging into the scaffold. The length of individual tubes produced by wild-type (WT) *H. pylori* ranged from 76 to 547 nm, with a median of 193 nm (n = 18). To our surprise, the tubes were not associated with obvious basal body-like densities localized directly beneath the tube in the periplasm or the inner or outer membranes, suggesting that a dedicated membrane-bound apparatus is either not required for tube formation, not recognizable at the current resolution, or had disassembled prior to sample freezing.

**Figure 1.**
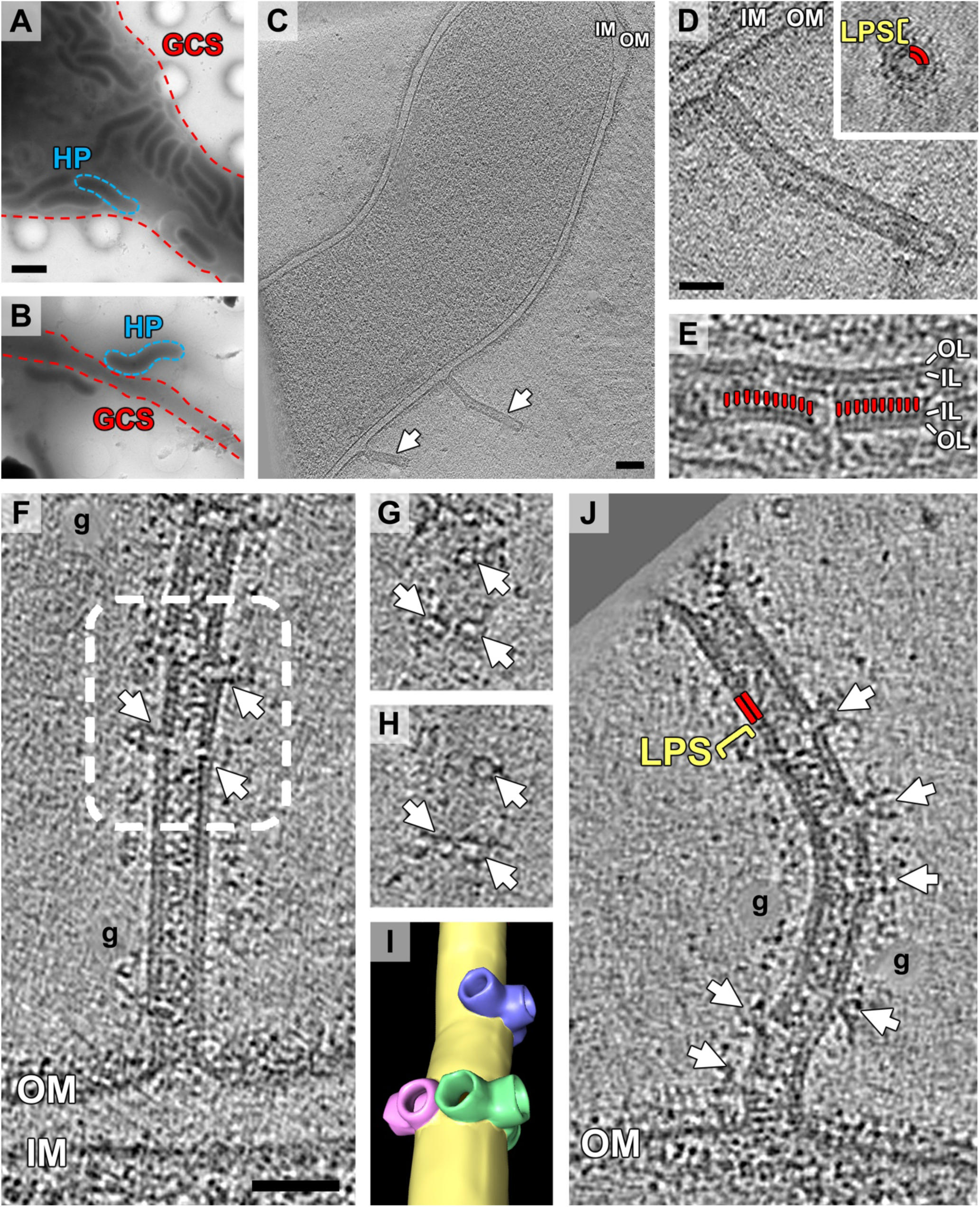
*H. pylori* assembles membranous tubes when in proximity to gastric epithelial cells. (A and B) Low magnification views of gastric epithelial cells grown on electron microscopy grids and infected with *H. pylori*. Blue dashed lines indicate examples of adherent *H. pylori* cells (HP) interacting with gastric cell surfaces (GCS; Red dashed lines) that were imaged by ECT. (C) Tomographic slice through WT *H. pylori* cell co-cultured with gastric epithelial cells. White arrows point to membrane tubes extending from the *H. pylori* cell envelope. (D) Enlarged view of the longer tube shown in C. Inset, cross-section of a membrane tube. The two leaflets of the tube’s membrane bilayer are labeled by two red lines; the region of lipopolysaccharide (LPS) is indicated by a yellow bracket. (E) Periodic densities lining the inside of the tube (labeled by red lines). (F) An individual tube displaying lateral ports (white arrows). (G and H) Distal and proximal cross sections of ports within the boxed region of the tube depicted in F. (I) 3D segmentation of the boxed region of the tube depicted in F. (J) The presence of lateral ports appears to induce a slight bending of the tube. The two leaflets of the tube and lipopolysaccharide densities decorating the surface are labeled as in D. The locations of erased gold fiducials during tomogram reconstruction are labeled with “g”. Scale bar in (A), 2 µm, applies to (A) and (B); (C) 100 nm; (D) 50 nm; (F) 50 nm, applies to (E-H) and (J). OM, outer membrane; IM, inner membrane; OL, outer leaflet; IL, inner leaflet.

To investigate whether the tubes were related to *cag* T4SS activity, we used ECT to image *H. pylori* lacking either the effector protein CagA, the *cag* T4SS pilus regulating protein CagH (*22*), or the entire *cag*PAI. ECT revealed tubes extending from the bacterial envelope when either *cagA* or *cagH* mutants were co-cultured with gastric cells (Fig. S2), but not the *cag*PAI strain. Under these conditions, tubes were produced by *H. pylori* proximal to a gastric cell in a total of 17 of 336 tomograms across the analyzed strains. We visualized roughly equivalent numbers of tubes per cell produced by WT, *cagA*, and *cagH* strains (*cagA*, n = 22; *cagH*, n = 23). Two extremely long tubes (785 nm and 1311 nm) were observed in the *cagH* mutant (Fig. S3 shows the longest observed tube), and this strain also assembled a few tubes with larger outer and inner diameter dimensions (Fig. S2).

### *In vivo* ultrastructure of the *cag* T4SS

In some tomograms, we noticed dense, periplasmic, cone-shaped particles spanning the bacterial envelope near (but not directly below) membrane tubes (Fig. 2A, B). These structures were reminiscent of *L. pneumophila dot/icm* T4SS complexes observed *in situ* (*19*), and consisted of distinct layers of densities in the periplasmic space near the outer membrane (Fig. S4A, B). Based on the structural similarity to the *dot/icm* T4SS, we hypothesized that these structures corresponded to a T4SS. Close inspection of our tomograms revealed varying numbers of these particles in each *H. pylori* cell (ranging from 0 – 4 particles per cell in the field of view of the tomograms). The particles were found at cell poles as well as sides, consistent with previous reports analyzing *cag* T4SS components (*21-24*). In many instances, we observed the bacterial outer membrane bulging to accommodate the assembled particle. We also captured several top views of the structure which revealed two concentric rings. The outer ring exhibited 14-fold symmetry and a diameter of 40 nm (Fig. 2C), consistent with the structure of immunopurified *cag* T4SS core complexes resolved earlier by negative stain electron microscopy (Fig. 4D) (*20*). Among the four potential T4SSs harbored by *H. pylori*, the imaged strain lacks complete *tfs3* and *tfs4* systems (*8*), and although the strain harbors the *comB* T4SS, corresponding cone-shaped particles were never observed in over 100 tomograms of the cognate *cag*PAI strain co-cultured with gastric epithelial cells, leading us to conclude that these particles are the *cag* T4SS rather than the *comB* DNA uptake or other T4SSs.

**Figure 2.**
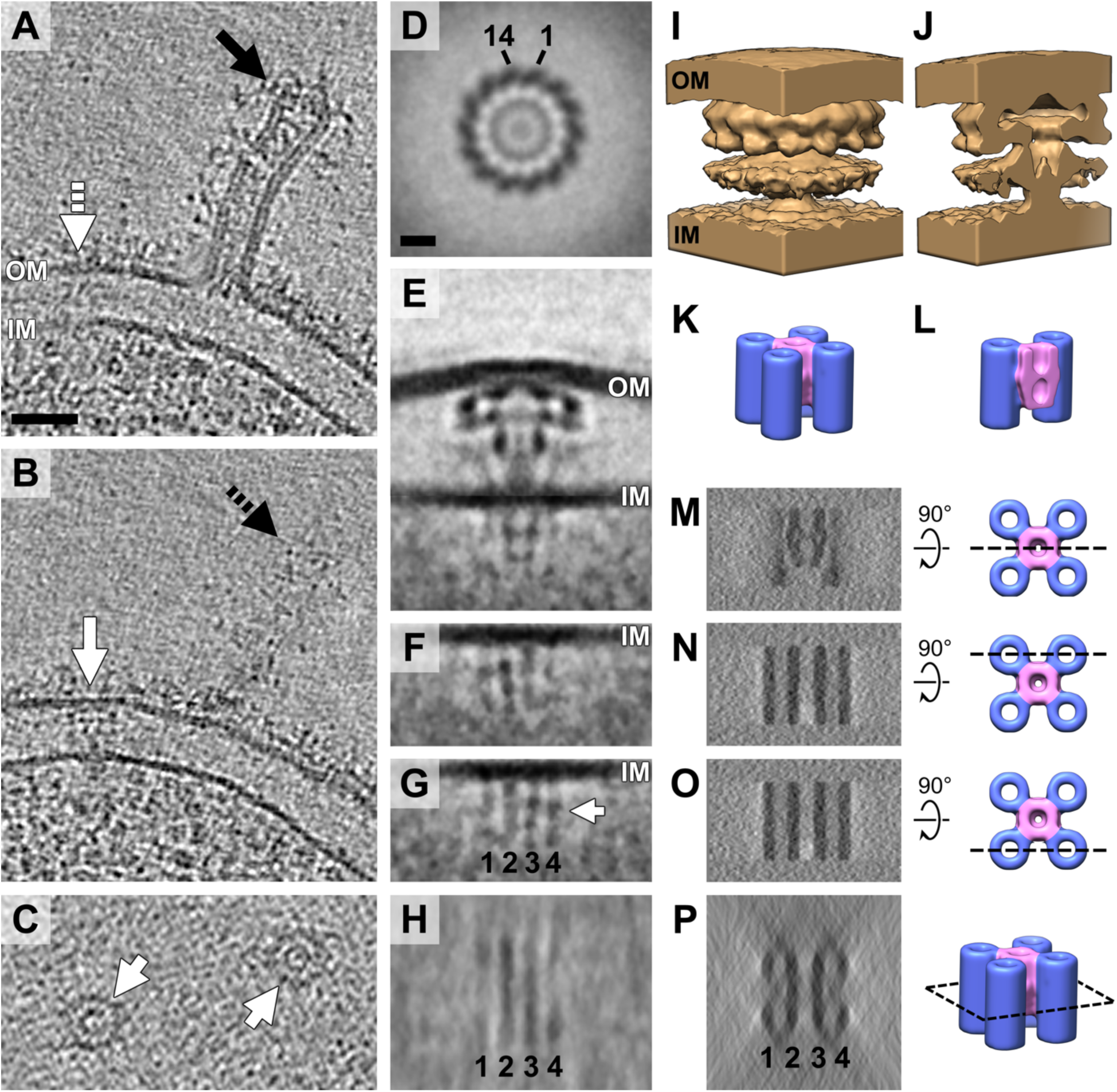
*In vivo* structure of the *cag* T4SS. (A and B) Different tomographic slices of the same region of the bacterial envelope identifying a *cag* T4SS particle (white arrow) adjacent to a tube-like appendage (black arrow). Dashed arrows represent the position of the tube and *cag* T4SS particle in the other tomographic slice. (C) Top views of individual *cag* T4SS particles (white arrows). (D) Top view of the subtomogram average of *cag* T4SS reveals 14-fold symmetry. Numbers indicate the clockwise count of individual subunits visible in the ring structure. (E) Central slice through the side view of the composite subtomogram average of the *cag* T4SS.Averages aligned on the periplasmic and cytoplasmic parts are stitched using the inner membrane as the boundary. (F) Distal and (G) proximal off-center tomographic slices of the cytoplasmic apparatus from the side view reveal four distinct rod-like densities. (H) A top view of the cytoplasmic apparatus at the level of the white arrow in G shows two central lines and four corner densities. (I, J) 3D representation (I) and cut-away view (J) of the *cag* T4SS periplasmic structure. (K, L) 3D representation (K) and cut-away view (L) of the predicted five-barrel structure of the cytoplasmic apparatus. The shorter central barrel is colored light pink. (M-P) Simulated tomograms of the five-barrel model of the cytoplasmic apparatus corresponding to tomographic slices E (M), F (N), G (O), and H (P). The position of each predicted tomographic slice is indicated in the views of the five-barrel model to the right. Scale bar in (A) 50 nm, applies to (A-C); (D) 10 nm, applies to (D-H). OM, outer membrane; IM, inner membrane.

To investigate structural details of the *cag* T4SS, we sought to generate a subtomogram average. In initial averages, we were able to resolve clear structural features in the periplasm but not the cytoplasm. Given the inherent structural flexibility of other T4SSs (*15, 19*), we aligned and averaged the periplasmic and cytoplasmic regions separately and then generated a composite average (Fig. 2E; Fig. S4C). In the periplasm, we resolved a “hat” density associated with the outer membrane, several ring-like densities surrounding and beneath the hat, a central stalk, and wing-like densities on the periphery (Fig. S4C, E). Cross sections through the cytoplasmic apparatus revealed parallel lines of density (Fig. 2E-H), but because most of the *cag* T4SS particles used in the average were imaged from the side (electron beam parallel to the membranes), the average was smeared by the missing wedge effect in that direction. To interpret these densities, we therefore explored a variety of candidate structures by generating artificial tomograms smeared by the same missing wedge effect, and then compared their cross sections to the experimental data (Fig. S5). We tested configurations of one to six barrel densities and various combinations of barrel and rod structures. The best-matching model consisted of a short central barrel surrounded by four longer barrels, which together recapitulated the experimental results very well (Figs. 2E-H, 2K-P, S5).

### Sheathed cytoplasmic rod on the *cag* T4SS

In three unusual *cag* T4SS particles (out of a total of 70 particles), we observed a dense, central rod extending from the outer membrane-associated complex into inner membrane invaginations with different depths (Fig. 3B-D, F-H). This feature was not observed previously in the *dot/icm* T4SS, nor in the purified R388 T4SS particles (*15, 19*). The rod and inner membrane invaginations measured ∼10 nm and 30 nm in width, respectively, and the rod extended 45 – 120 nm from the outer membrane complex (Fig. 3B-D). Notably, in one of the particles, the rod appeared to project through the inner membrane into the cytoplasm, though the details were obscured by the crowded cytoplasm (Fig. 3D, H).

**Figure 3.**
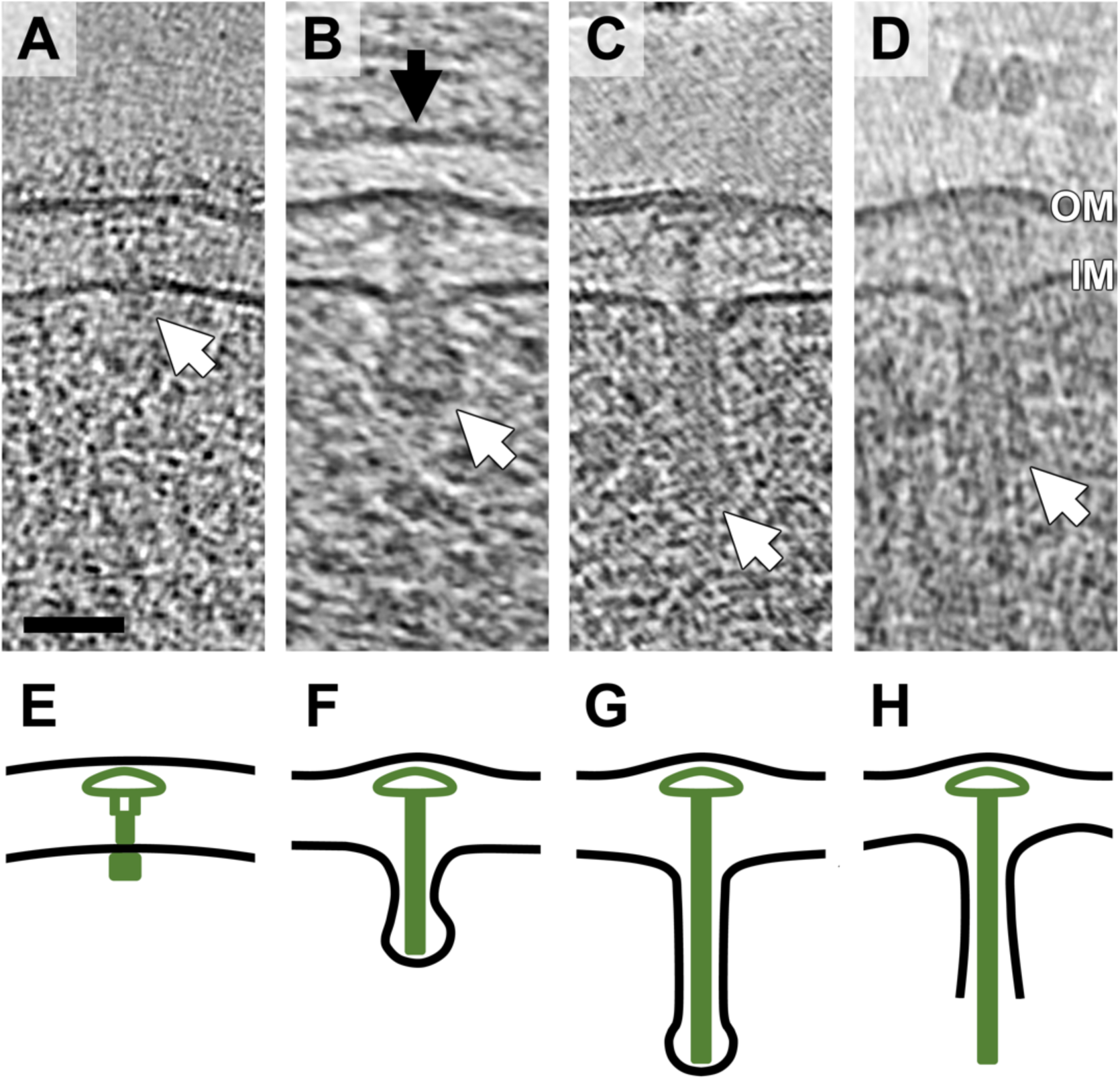
Pilus-like rods emerging from the *cag* T4SS. (A) A *cag* T4SS particle exhibiting a typical cytoplasmic structure (white arrow). (B) A *cag* T4SS particle with a pilus-like rod density surrounded by an inner membrane invagination (white arrow). The gastric epithelial cell plasma membrane (black arrow) is visible directly above the bacterial outer membrane. (C) A *cag* T4SS particle with an extended pilus-like rod density and inner membrane sheath (white arrow). (D) A *cag* T4SS particle with an even longer pilus-like rod density. The inner membrane appears to have ruptured (white arrow). (E-H) Schematic interpretation of the *cag* T4SS apparatus conformation in A (E), B (F), C (G), and D (F). Scale bar in (A) 50 nm, applies to (A-D). OM, outer membrane; IM, inner membrane.

### Comparison to previous T4SS structures

The T4SS family is phylogenetically diverse, and has been divided into two major sub-types, type IVA and type IVB (T4ASS and T4BSS). Historically, T4ASSs have been classified according to protein homology to components of *E. coli tra* DNA conjugation systems (types F and P) and the *A. tumefaciens vir* T4SS, while T4BSSs exhibit protein sequence conservation to IncI-like conjugation systems and the *L. pneumophila dot/icm* T4SS (*28*). In most cases, T4ASSs are comprised of approximately 12 components with clear homology to Vir proteins, while T4BSSs incorporate many more proteins (twenty or more), and few share sequence homology with *vir* T4SS components (*28*). The *cag* T4SS has been considered a T4ASS since several Cag proteins share limited sequence similarity to Vir components (*6, 28*); however, the homologies are so weak their relevance is unclear, and the *cag*PAI encodes as many genes as a typical T4BSS, including many *H. pylori*-specific genes (*6*). Thus, the *cag* T4SS may represent a mosaic or hybrid T4SS subtype.

To explore the structural relationships between T4SSs, we compared the *cag* T4SS sub-tomogram average to the previous EM and crystallographic structures of purified sub-complexes of the R388 (*15*) (Fig. 4A, B), EM images of negatively-stained immuno-purified subcomplexes of the *cag* T4SS (Fig. 4C, D), and the subtomogram average of the *in vivo dot/icm* T4SS (*19*) (Fig. 4E; Fig. S4E). In comparison to the R388 structures, the *cag* T4SS is similar in that it includes a large cluster of densities associated with the outer membrane, a stalk that connects the outer membrane-associated cap to the inner membrane, and structures in the cytoplasm anchored to the inner membrane (Fig 4A, G). The size and shape of the R388 outer-membrane-associated cluster (referred to in (*16*) as the “core” complex, containing the C-termini of VirB7, VirB9, and VirB10) matched the hat and inner ring density below the hat (labelled δ in Fig. S4E) in the *cag* T4SS structure (Fig. 4A, B, G, H, orange demarcation). Because CagY shares low-level homology to VirB10 (*6, 21*) and CagX shares low-level homology to VirB9 (*6, 29*) in those same C-terminal regions, we reason that these regions of CagX and CagY form the hat and the density labeled δ. The size of the stalk and the configuration of the cytoplasmic barrels in the two structures, however, appear quite different.

**Figure 4.**
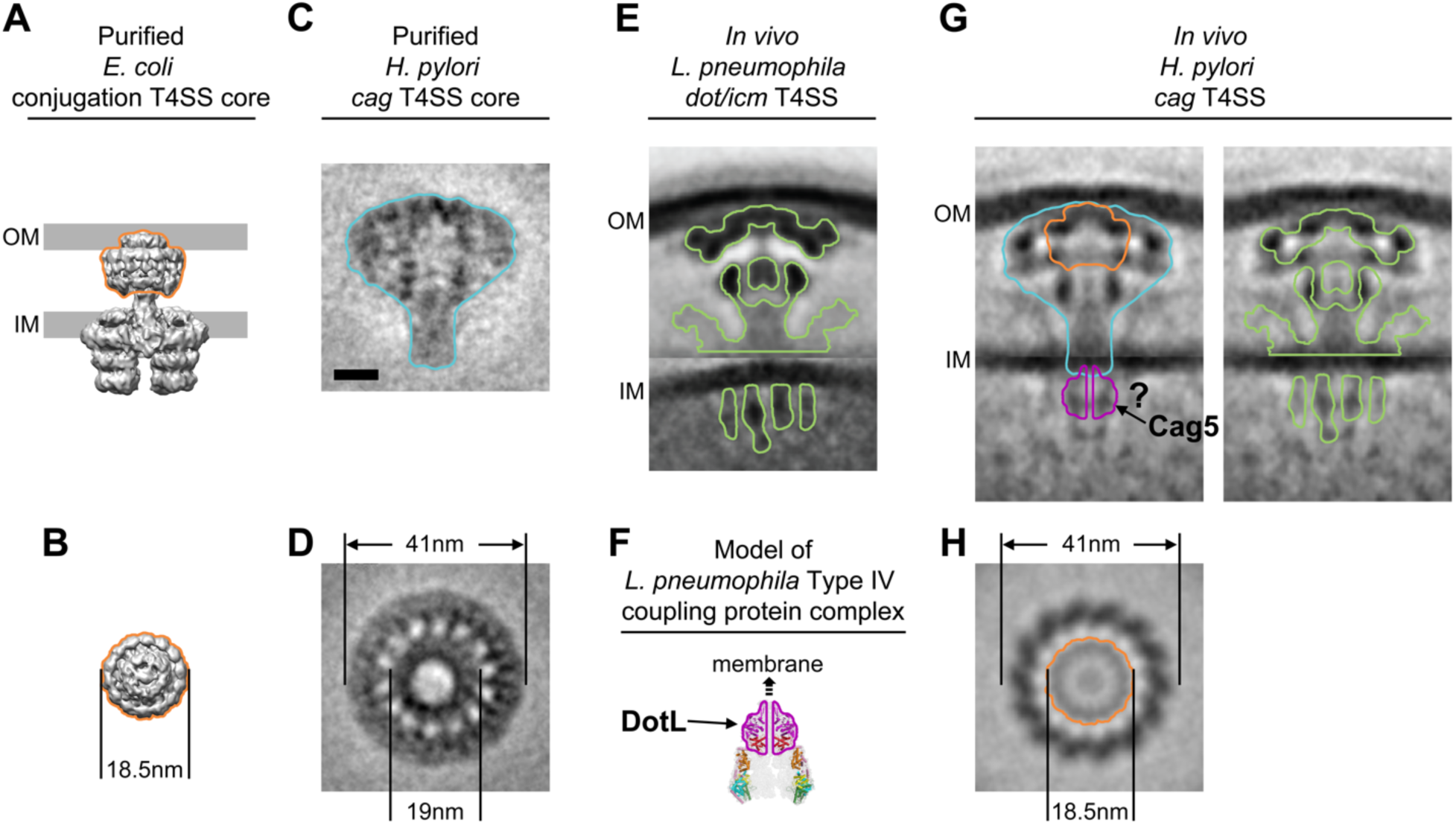
Comparison of diverse T4SS machinery structures. (A) Side and (B) top views of the purified *E. coli* R388 conjugation system (reproduced from (*15*)). (C) Side and (D) top view of immunopurified *cag* T4SS core complex particles (adapted and modified from (*20*)). (E) Side view of the *L. pneumophila dot/icm* T4SS *in vivo* (*19*). (F) Model of the *L. pneumophila* DotL coupling protein complex (*30*), with the DotL structure outlined in magenta. (G) Side and (H) top view of the *cag* T4SS structure *in vivo* (present study). In (G), two duplicated side views are shown for clearer labeling. Orange outline indicates the R388 core complex positioned within the *cag* T4SS structure; blue outline indicates the position of the purified *cag* T4SS core complex within the *in vivo* structure; magenta outline indicates the predicted location of the coupling protein Cag5 based on the structure of its *L. pneumophila* homolog, DotL, shown in (F); green outline indicates the *L. pneumophila dot/icm* T4SS structure superimposed on the *cag* T4SS structure. Scale bar in (C) 10 nm, applies to all panels. See also Fig. S4 and S6.

In comparison to the images of the purified *cag* T4SS sub-complex comprised of CagM, CagT, CagX/VirB9, CagY/VirB10, and Cag3 (Fig. 4C, G, blue demarcation), the periplasmic portion of the *in vivo cag* T4SS average exhibited almost the exact size and general shape, which allowed us to definitively position and orient the negative stain result relative to the bacterial envelope. Comparison of the particle top views (Fig. 4D, H) also revealed striking structural similarities, including in the sizes of the concentric rings and their 14-fold symmetry (Fig. 4D, H) (note that the relative contrast of features in images of negatively stained proteins can depend on surface chemistry and stain accumulation, so the shapes and arrangement of densities are the important properties to compare rather than grey-levels). These observations further confirm that the particles averaged in this study are the *cag* T4SS. From this comparison, and based on previous experimental evidence (*20*), we conclude that CagT, Cag3, and CagM must produce the densities inside the blue outline but outside the orange in Figure 4G (which were already assigned to CagX/VirB9 and CagY/VirB10). CagT and Cag3 can be further pin-pointed as the upper and lower outer rings (α and β in Fig. S4E) based on the published negative-stain images of the CagX, CagY, and CagM sub-complex (*20*), which are missing those rings (though which is the upper and which is the lower ring or if they are mixed remains unclear). By elimination, this suggests CagM produces the density labeled γ in Fig. S4E; however, CagM localization could be more complicated and so remains to be verified.

Compared to the *in vivo dot/icm* T4SS structure, the *cag* T4SS structure is remarkably similar, considering the fact that the systems each comprise over 25 components and only a few share sequence homology (Fig. S6). Both structures exhibit (i) an outer-membrane associated hat; (ii) upper and lower outer ring-like densities surrounding the hat (labeled α and β in Fig. S4E, F); (iii) a barrel-like γ density at the center of the structure; (iv) a central stalk; (v) weak, wing-like densities that extend from the inner membrane into the periplasm (Fig. S4C-F), and (vi) parallel elongated densities perpendicular to the membrane in the cytoplasm. While the upper and outer ring densities (α) superimpose well (Fig. 4G, right panel), a difference is that there are two densities in the lower ring of the *cag* T4SS (labeled β and δ in Fig. S4E) versus only one (labelled β in Fig. S4F) in the *dot/icm* T4SS. Based on a recent report describing the structure of the *L. pneumophila dot/icm* coupling protein DotL (*30*), its predicted position within the secretion system just underneath the IM (*30*), and its match in size and shape to the central barrel seen at the same location in the *H. pylori* subtomogram average, we speculate that the central barrel of the *cag* T4SS cytoplasmic density corresponds to the Cag5/VirD4 coupling protein (Fig. 4F, G, magenta demarcation; Fig. S6). The tentative positioning of Cag5 to the central barrel density of the cytoplasmic apparatus is further supported by recent work demonstrating that the VirD4 coupling protein is situated in the center of the R388 inner membrane complex between VirB4 barrels (*31*). Collectively, these data reveal that although the *cag* T4SS is phylogenetically distinct from both the R388 and the *dot/icm* T4SS, the gross architecture of these three T4SS machines is remarkably conserved.

## Discussion

Here we have reported the structure of the *cag* T4SS and shown that when *H. pylori* are in proximity to host cells, the bacterium produces membranous tubes decorated with pipe-like ports. Multiple scanning electron microscopy (SEM) studies have shown that under similar conditions, *H. pylori* assembles extracellular filaments, but the drying and metal coating inherent to SEM obscured fine details (*21-23, 25, 26*). In previous papers, these structures have been referred to as “pili,” “*cag* T4SS-associated pili,” “filaments,” “extensions,” etc., but here we will refer to all of the previously observed structures as “SEM filaments” for clarity, and because we would like to use the word “pilus” for a single component of the structure (the rod) described below. We conclude that the membrane tubes visualized by ECT are the native form of the previously described SEM filaments for the following reasons. First, both the membrane tubes and SEM filaments were only seen when *H. pylori* was co-cultured with gastric epithelial cells. Second, in all cases both structures depended on the presence of the genes in the *cag*PAI. Third, one previous study interpreted the SEM filaments as proteinaceous pili covered by membrane sheaths (*21*).

Fourth, and most decisively, deletion of the CagH “molecular ruler” protein resulted in both longer and wider membrane tubes and longer and wider SEM filaments (*22*). Concerning size, unfortunately the different SEM studies reported substantially different diameters for the SEM filaments, ranging from 15-70 nm (*21-25*). While this range does include the diameter of the native membrane tubes measured here (consistently 37 nm median diameter), we speculate that in the previous SEM studies, the extensive chemical fixation, dehydration, and metal coating inherent to the method may have introduced the variations.

Previous SEM immuno-labeling experiments showed that CagY is present along the SEM filaments (*21, 26*) and CagT is clustered in ring-like formations at the SEM filaments’ base (*21*). Other immuno-labeling studies showed that additional Cag proteins could be localized to the SEM filaments, including CagA, CagL, CagT, and CagX (*21, 23, 24, 26, 29*). Comparisons of the T4SS structure obtained here with previous structures and images of purified complexes revealed that the conserved C-terminal region of CagY forms the central part of the “hat” density and that CagT forms part of the outer ring. Assuming CagX is a homolog of VirB9 as predicted (*6, 29*), CagX is also present in the hat density. Because both the *cag* T4SS and SEM filaments/membrane tubes have been associated with CagY, CagT, and CagX, we propose that the *cag* T4SS and the tubes are different states of the same secretion apparatus. In support of this, we note that the outer and inner diameters of the membrane tubes (37- and 22-nm, respectively) approximately match the outer and inner ring dimensions of the T4SS (41- and 19-nm, respectively).

More specifically, we propose as a working hypothesis that the T4SS structure shown here (Fig. 2E) is a “pre-extension” state that assembles in response to contact with a host cell (Fig. 5A). It is known that the *E. coli tra* and *A. tumefaciens vir* T4SSs produce protein pili (Fig. 5B). In the *tra* system, the protein F pilus has an outer diameter of 8.7 nm and is formed by the major pilin TraA/VirB2, which is otherwise found in the inner membrane (*32*). The images shown in Fig. 3 reveal that the *cag* T4SS also produces a rod-like structure with similar diameter, seen here in the periplasm. Upon receiving the proper signals, we therefore propose that the *cag* T4SS also assembles a protein pilus (the rod) from subunits in the inner membrane. This pilus may be formed of CagC, which exhibits weak homology to the VirB2/TraA component in the *vir* T4SS (*25, 33, 34*). In our working model, we propose that as the pilus grows upwards from the IM, it interacts with components of the core complex within the periplasm, possibly opening a translocation channel through the system (*1*). At the onset of membrane tube formation, a conformational change within the core complex disengages the CagX/CagY hat which is released from the CagT/Cag3 outer ring and extended tubes are produced by growth of the thin protein scaffold lining the inside tube surface (Fig. 5C). The tubes are stabilized by the CagT/Cag3 outer ring, which remains at the base like a collar, and the scaffold, which holds their diameter constant along their length. This scaffold may contain CagL, which has been proposed to be a functional homolog of the VirB5 subunit (*35*) that is known to decorate the outside of the VirB2 T-pilus in *A. tumefaciens* (*36, 37*). If some part of CagL extended through the outer leaflet of the tube lipid bilayer, it would explain the observations that CagL can bind host cell integrins (*23*) and can be localized to the SEM filaments by immunogold labeling (*23*). CagI, which can also bind integrins (*26*) may also be part of the scaffold (*22, 38*). This model would explain why CagL and CagI deletion mutants have no SEM filaments (because tubes do not form in the absence of the scaffold) (*22*). Given that CagH is membrane-bound, plays an essential role in T4SS activity, regulates SEM filament/tube dimensions, and forms a complex with CagL and CagI (*22*), CagH may control the incorporation of CagI and CagL into the scaffold (*22, 38*).

**Figure 5.**
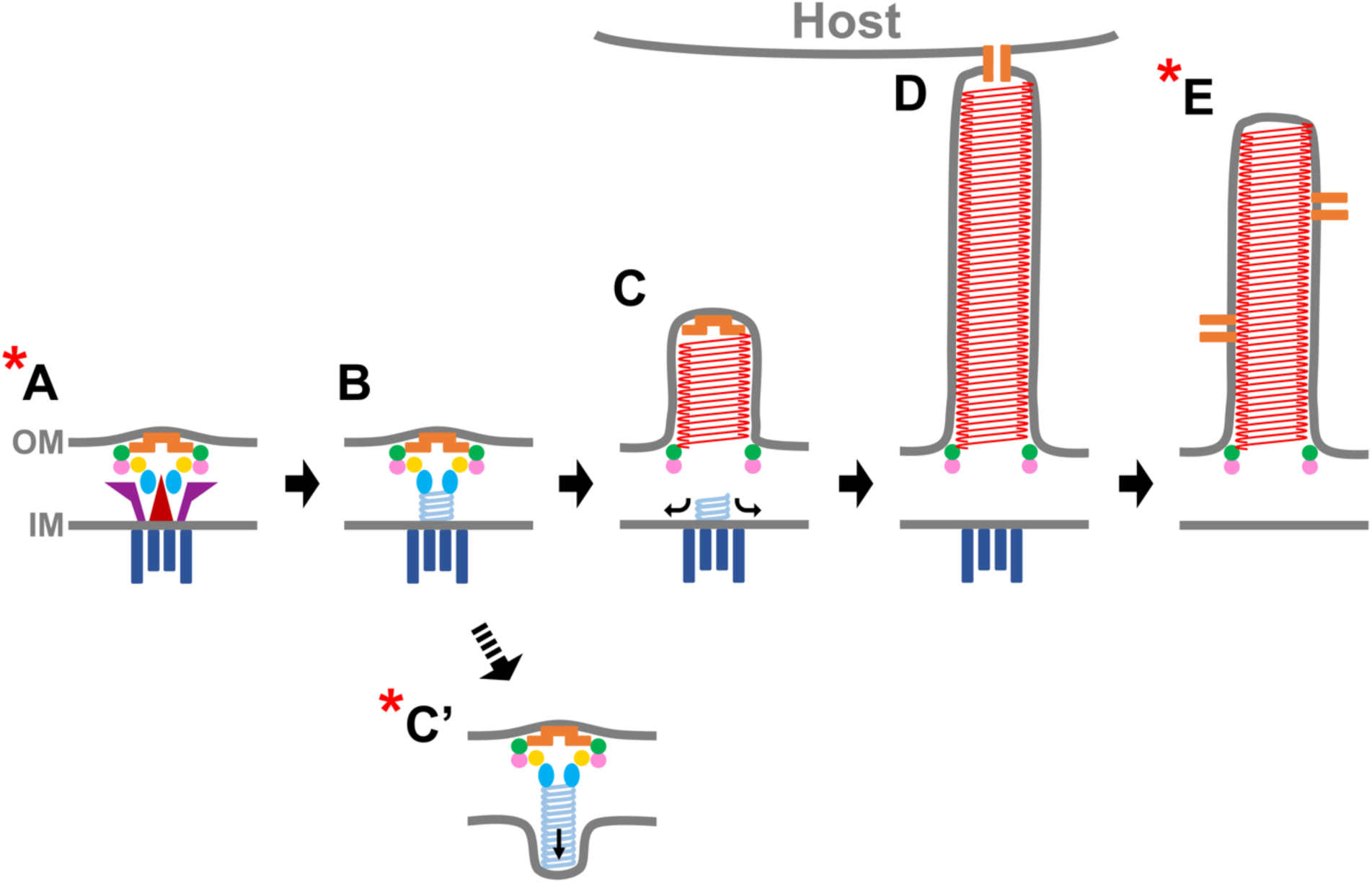
Working model for biogenesis of the *cag* T4SS. See Discussion for proposed identities and roles of structures. The starred structures (A, C’, and E) are the long-lived states observed in the cryotomograms, while other stages of assembly are hypothesized.

We propose that the CagX/CagY hat is a membrane fusion machine positioned at the tip of the tubes. Upon contact with a host cell, it fuses the tube and host membranes, opening a channel for the passage of effectors (Fig. 5D). Our interpretation of the tube ports is that they are open CagX/CagY channels, and this may explain the immuno-labeling results that CagY can be localized along the length of the SEM filaments (*21, 26*), and the observation that the C-terminal/VirB10 domain region of CagY can bind host integrins (*26*). While one CagX/CagY complex is positioned at the tip of the tube, other complexes present in the OM may be drawn upwards into the tube as it extends, or alternatively, additional CagX/CagY complexes may assemble in the tubes’ lateral walls after extension. The interpretation that CagY is the membrane fusion protein could explain why strains lacking *cagY* form SEM filaments, but do not secrete effector molecules (*24, 25*), and is consistent with the proposal that CagY serves as a molecular switch that regulates secretion activity *in vivo* (*24*). In the imaged strains, the lateral ports exhibited a diameter of ∼10-12 nm, which is large enough to facilitate transport of the folded CagA effector protein, whose dimensions measure 8 by 11 by 5.5 nm (*39*). One problem with this model is what happens to the IM transmembrane domain of CagY – given its tether to the IM, how would it remain at the tip of the extending tube? One possibility is that as the IM is perturbed by pilin subunits being loaded out of the IM and onto the base of the pilus, and as the pilus itself may incorporate IM lipids similar to *E. coli tra* F pili (*32*), the CagY tether is somehow released. Another possibility is that as CagY is ∼1900 amino acids long, it spans the length of the tube.

Our interpretation of the three unusual T4SS particles with pili protruding downwards towards the cytoplasm (Fig. 3) is that these are stalled end-states in which the CagX/CagY hat failed to disengage the CagT/Cag3 ring, forcing pilus growth to push the IM downwards instead of the OM upwards. It may therefore be that the only states captured in cryotomograms are long-lived, including the pre-extension state (Fig. 5A), various stalled “failure to fire” states (Fig. 5C’), and terminal end states of tubes no longer connected to the host, leaving the tubes to reseal at the tip (Fig. 5E). Future studies will focus on earlier stages of the association and on bacteria directly touching host cells, in hopes of producing images of the hypothesized transitory states (Fig. 5B-D).

The role and fate of the pilus itself remains particularly unclear. While pilus growth might be involved in tube extension, a recent study revealed that SEM filaments can be produced by a strain lacking *cagC* (*25*), and it is also known that strains lacking *cagC* do not secrete *cag* T4SS effectors (*25, 34*). Perhaps there is some required interaction between the putative CagC pilus and the CagX/CagY hat that primes the hat for membrane fusion. Alternatively, as proposed for VirB2 in other type IVA systems (*1*), CagC may form a stable, rod-like translocation channel or pore through the periplasmic core complex into the base of the tube when the system is actively secreting effector molecules.

Assuming our model is correct, the *H. pylori cag* T4SS differs from the *E. coli tra* or *A. tumefaciens vir* systems in that the *cag* T4SS produces an extracellular appendage enclosed by outer membrane. Perhaps all T4SSs will share the basic central machinery that loads a VirB9/VirB10 membrane-fusion machine at the tip of an appendage (pilus or tube) that then extends to open a channel into a host cell, but differ in the presence and roles of peripheral proteins that manage the OM’s involvement in that appendage. Functionally, exposed protein pili may alone be sufficient to translocate single-stranded DNA (*40*), but wide membrane tubes like those seen here are likely required to translocate folded effector proteins into a host. Interestingly, some have already also proposed that membrane tubes are involved in DNA translocation as well (*41, 42*).

## Experimental Procedures

### Bacterial strains and growth conditions

*H. pylori* strain 26695 and corresponding mutants (*22*) were routinely maintained on Trypticase soy agar supplemented with 5% sheep blood (BD Biosciences) under microaerobic conditions. For all experiments, *H. pylori* were seeded into Brucella broth supplemented with 10% fetal bovine serum (FBS), and were grown overnight in shaking culture at 37°C, 5% CO_2_. Overnight bacterial cultures were normalized to an optical density at 600 nm (OD_600_) to ∼0.3 in fresh Brucella broth supplemented with 10% FBS, and cells were grown to mid-log phase at 37°C, 5% CO_2_ prior to generating samples for microscopy analysis.

### Human cell culture

The gastric adenocarcinoma cell line AGS (ATCC CRL-1739) was maintained in RPMI 1640 medium supplemented with 10% FBS, 2 mM L-glutamine, and 10 mM HEPES buffer (complete RPMI). Cells were grown at 37°C in 5% CO_2_.

### Sample preparation for electron cryotomography

AGS cells were seeded onto freshly glow discharged, UV sterilized Quantifoil gold Finder holey carbon grids (Quantifoil Micro Tools). AGS cells were added dropwise to the surface of the grid, and cells were grown overnight at 37°C in 5% CO_2_ to permit adherence to the grid. Grids were screened for cell confluency, and grids containing AGS cell clusters were inoculated with *H. pylori* at a multiplicity of infection (MOI) of approximately 50 bacterial cells per AGS cell. Co-culture samples were incubated in complete RPMI at 37°C in 5% CO_2_ for 4.5 h prior to mixing with 20 nm colloidal gold beads (Sigma-Aldrich) that were pre-coated with bovine serum albumin. Grids were blotted and plunge-frozen in a liquid ethane/propane mixture (*43*) using an FEI Vitrobot Mark IV (Thermo Fisher Scientific), and were stored in liquid nitrogen prior to imaging.

### Electron cryotomography data collection and processing

Frozen-hydrated samples were imaged in an FEI Polara 300 keV field emission gun transmission electron microscope (Thermo Fisher Scientific) equipped with a Gatan energy filter and a Gatan K2 Summit direct detector. Energy-filtered tilt-series of images of the cells were automatically collected from −60° to +60° at 1° intervals using UCSF tomography data collection software (*44*), with a total dosage of 160 e–Å^-2^ per tilt-series, a defocus of −6 μm and a pixel size of 3.9 Å. The images were aligned using the IMOD software package (*45*). SIRT reconstructions were then produced using the TOMO3D program (*46*). The *cag* T4SS structures were located by visual inspection. The sub-tomogram averages were produced using the PEET program (*47*) with local masks on either the periplasmic or cytoplasmic portion.

## Data availability

The sub-tomogram averages of *cag* T4SS machinery that support the findings of this study have been deposited in the Electron Microscopy Data Bank (EMDB) with accession codes EMD-XXXX (aligned on the periplasmic region); EMD-XXXX (aligned on the cytoplasmic region).

## Acknowledgements

We thank Dr. Tim Cover (Vanderbilt) for providing *H. pylori* strains and Dr. Maria Hadjifrangiskou (Vanderbilt) for helpful discussions. This work was supported by NIH grant R01 AI127401 to G.J.J., NIH grant F32 DK105720 to C.L.S., NIH grant 5 T32 A1007474-19 to C.L.S, and the Burroughs Wellcome Fund 2016 Collaborative Research Travel Grant to C.L.S.

## Author contributions

Y.-W.C, C.L.S. and L.A.R. prepared the samples. Y.-W.C collected and processed the electron cryotomography data and generated the subtomogram averages. Y.-W.C, C.L.S., L.A.R. and D.G. analyzed the electron cryotomography data. G.J.J supervised the project. Y.-W.C., C.L.S. and G.J.J. wrote the paper with input from all authors.

---

**Supplementary Figure S1.**
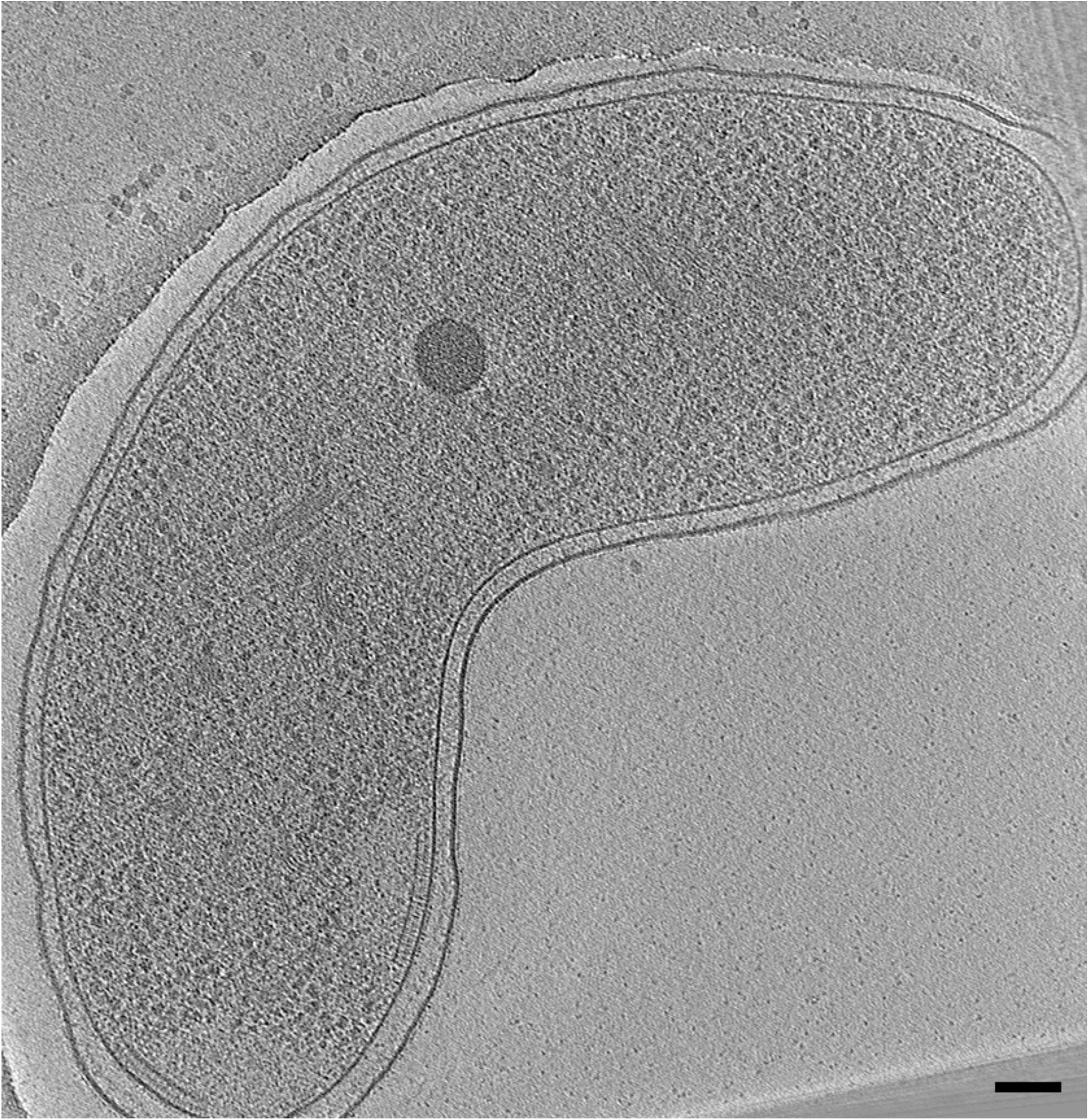
A tomographic slice of *H. pylori* without co-cultured with human gastric epithelial cells. Scale bar 100 nm.

**Supplementary Figure S2.**
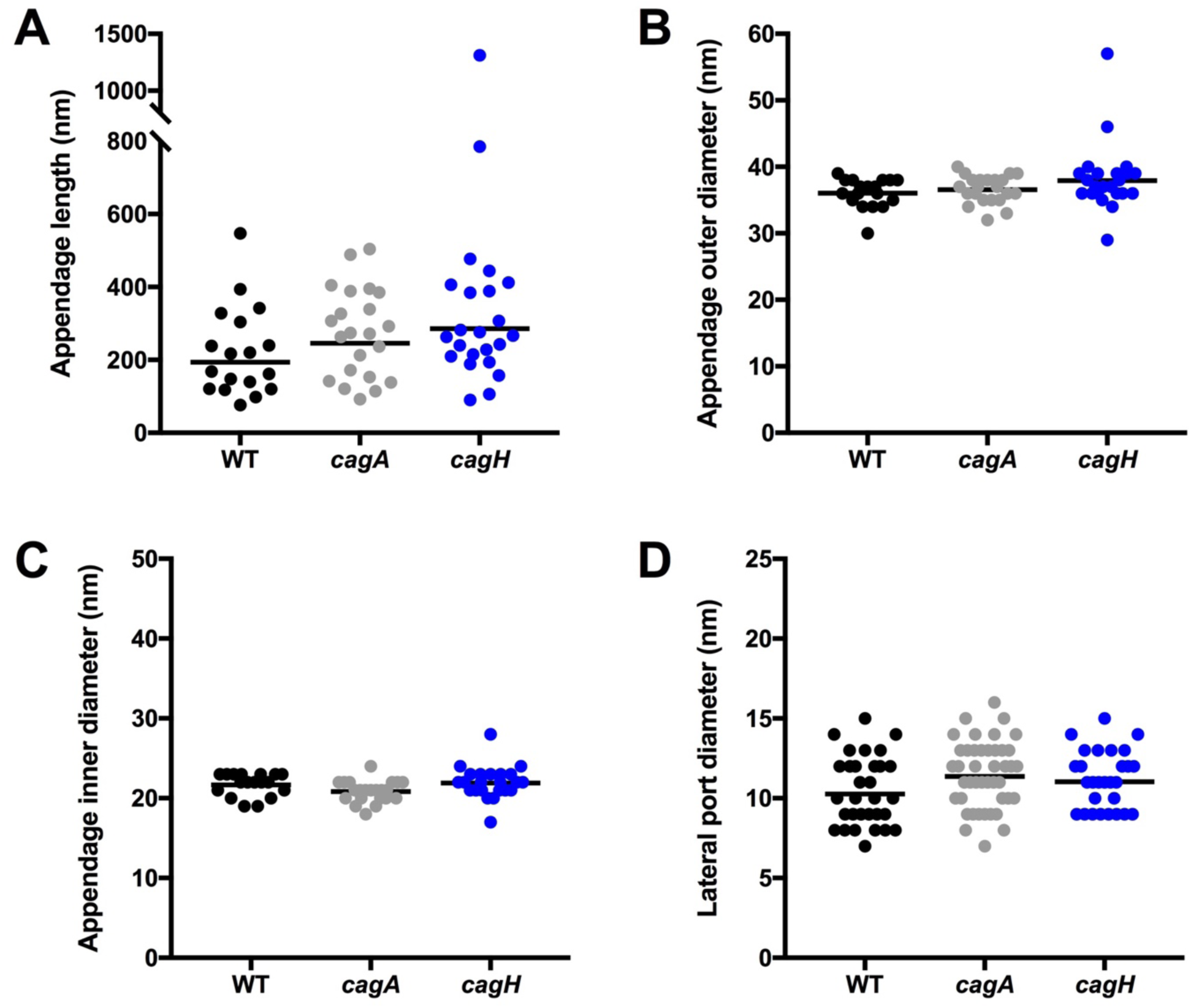
Dimensions of *H. pylori* membrane tubes. (A) Tube length for each imaged strain. (B) Outer and (C) inner diameter of bacterial tube structures. (D) Width of lateral pipe-like conduits associated with some tubes. Dots in (A-C) represent the dimensions of individual tubes; dots in (D) represent the diameters of individual lateral ports. Lines represent the geometric mean of each distribution.

**Supplementary Figure S3.**
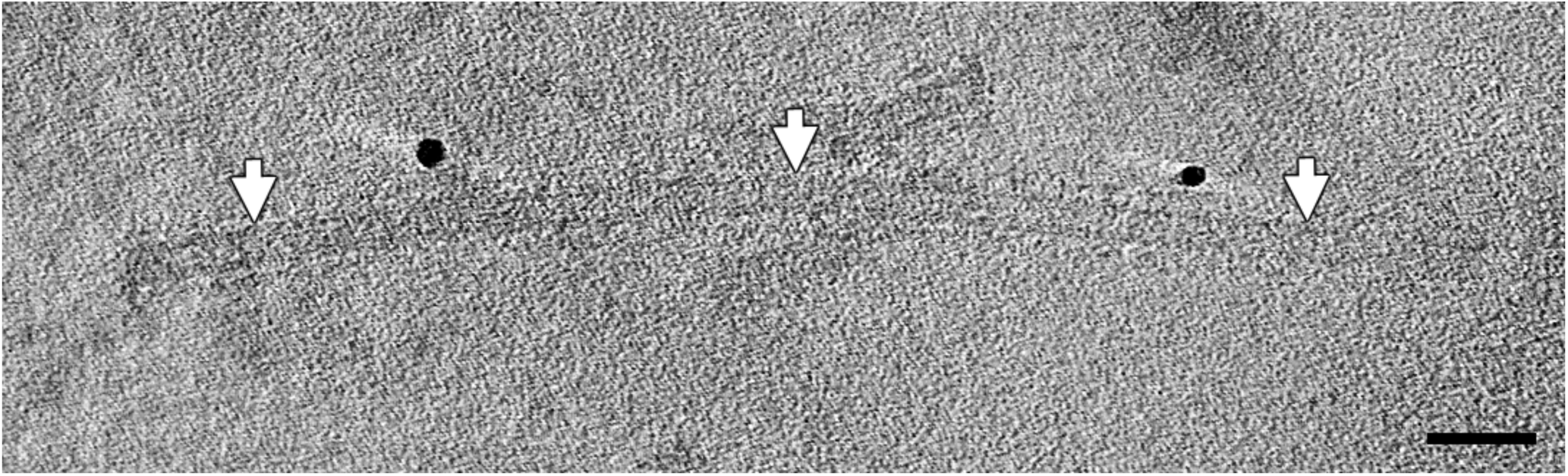
A tomographic slice of a >1,300 nm long tube (arrows) captured on the *H. pylori cagH* knockout strain in co-culture with human gastric epithelial cells. Smaller adjacent tubes can be seen flanking the central, long tube. Scale bar 100 nm.

**Supplementary Figure S4.**
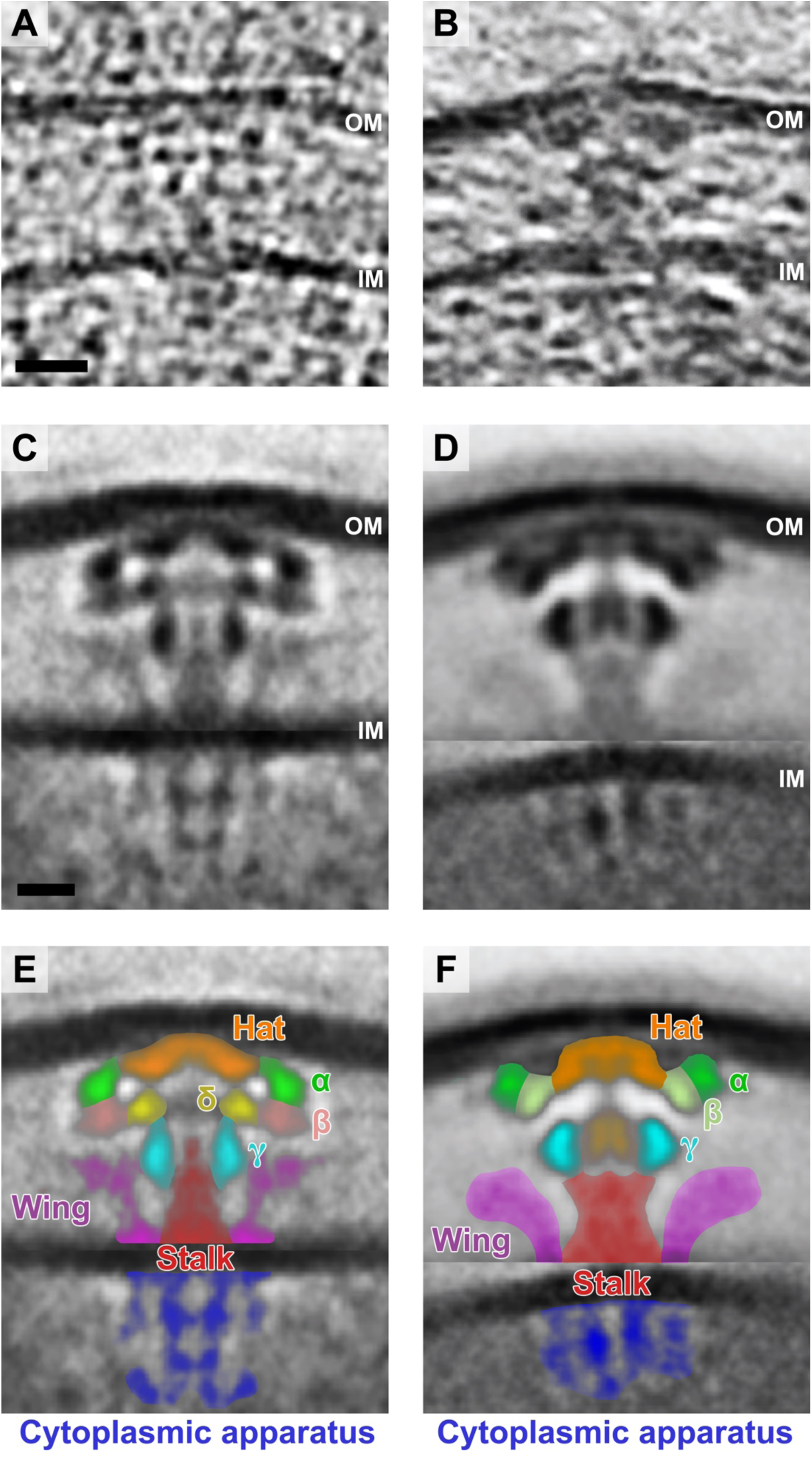
Schematic of structural features associated with the *cag* T4SS. (A) Cryotomographic slice 7.8 nm thick through an individual *cag* T4SS particle in a tomogram of *H. pylori*. (B) Cryotomographic slice 7.8 nm thick through a *dot/icm* T4SS particle in a tomogram of *L. pneumophila.* (C) Subtomogram averages of the *cag* T4SS (this study) and (D) the *L. pneumophila dot/icm* T4SS (*19*). (E and F) Colorized densities observed in subtomogram averages shown in C and D, respectively. Scale bar in (A) 20 nm, applies to (A, B); scale bar in (C) 10 nm, applies to (C-F). OM, outer membrane; IM, inner membrane.

**Supplementary Figure S5.**
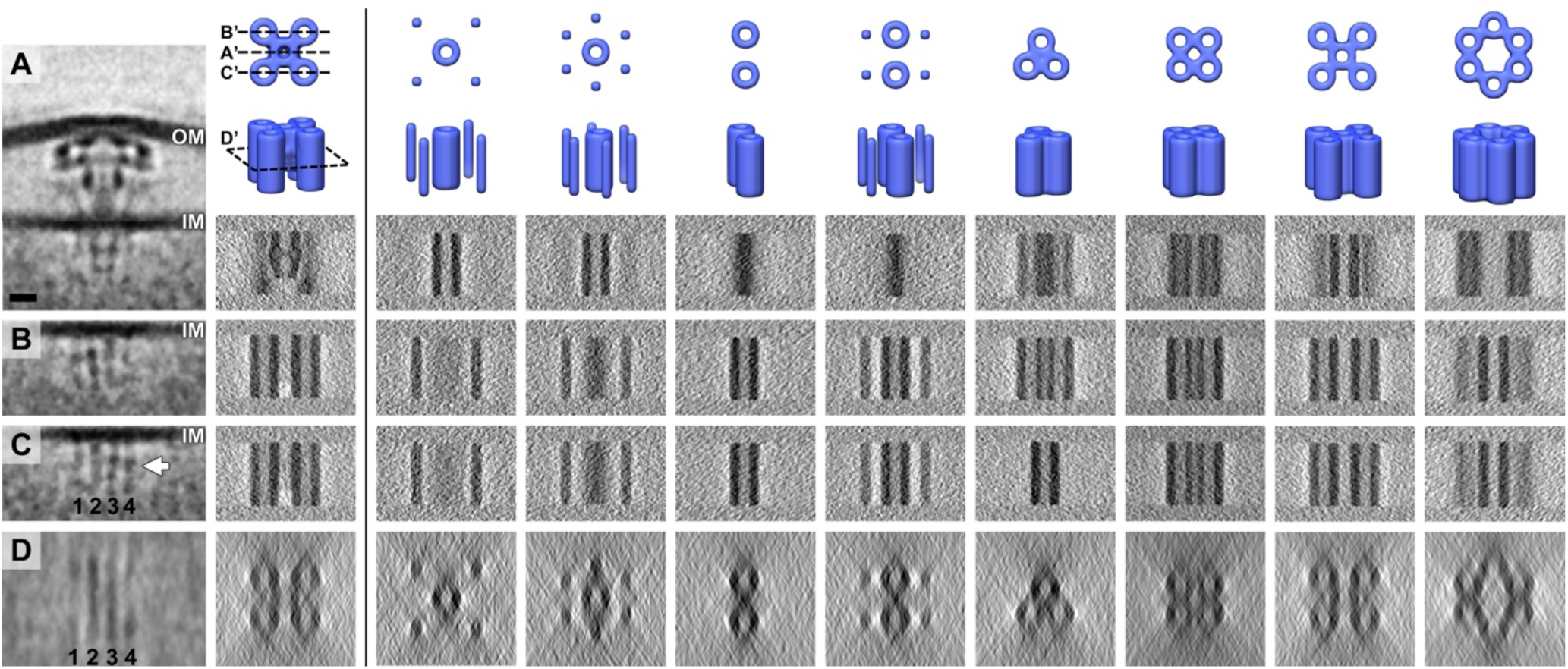
Modeling potential structures of the *cag* T4SS cytoplasmic apparatus. (A) Central slice of the subtomogram average of the *cag* T4SS from the *en face* view. (B) Distal and (C) proximal off-center tomographic slices through the *en face* view of the *cag* T4SS particle. (D) Cross-section of the cytoplasmic apparatus parallel to the inner membrane at the position indicated by the white arrow in (C). From the second column onward, the top two images are the top and side views of the candidate 3D models for the cytoplasmic apparatus. The four tomographic slices below the 3D models in each column are the sections corresponding to the first column (A-D). In the second column, A’, B’, C’ and D’ on the top two views of the 3D representation indicate the locations of the tomographic slices shown in (A-D) through the object. Comparing candidate structural models to the experimental data, we predict that the cytoplasmic apparatus is comprised of a five-barrel arrangement. Scale bar 10 nm, applies to all panels. OM, outer membrane; IM, inner membrane.

**Supplementary Figure S6.**
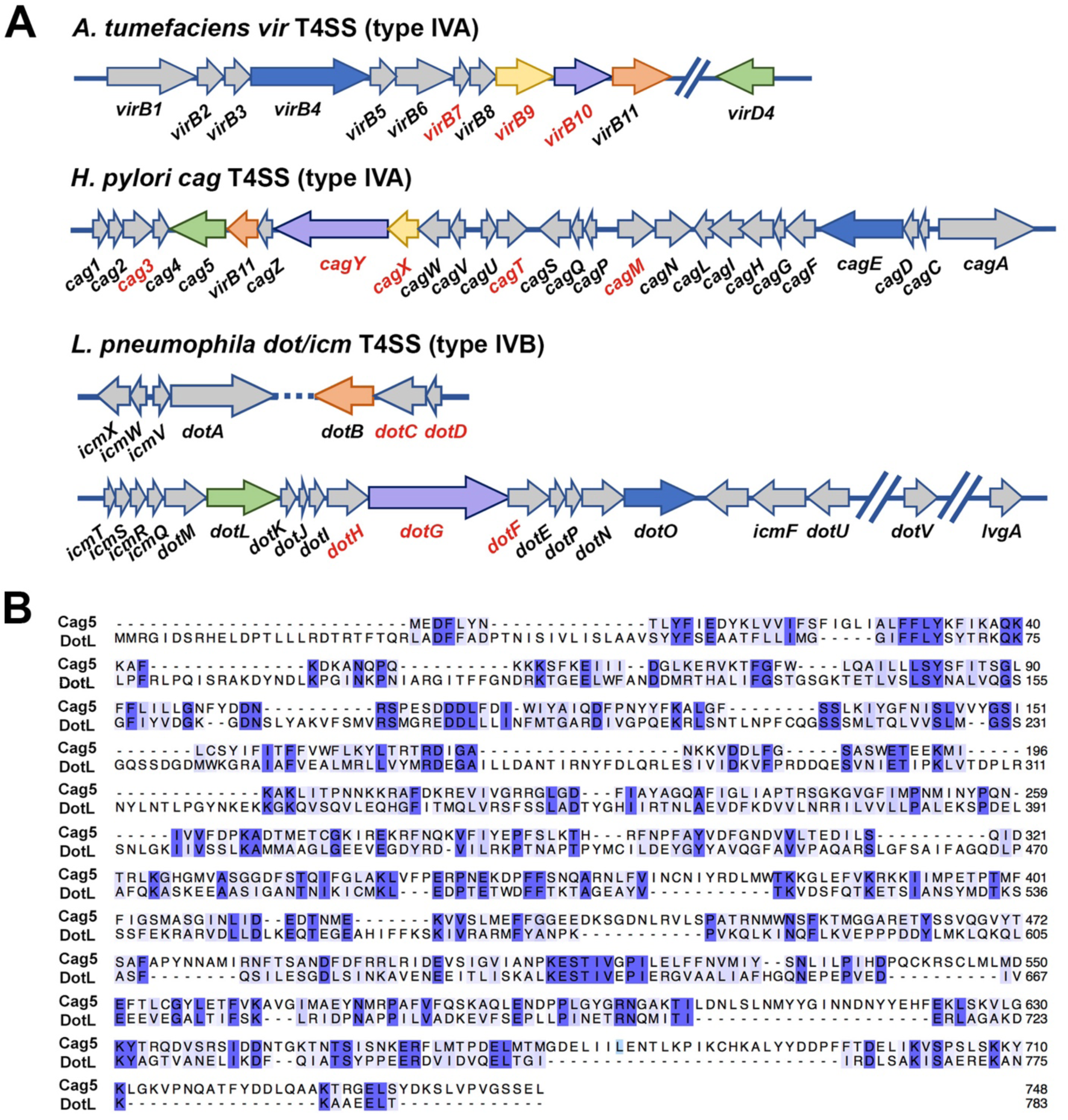
Genetic organization of T4SS-associated loci. (A) Organization of *A. tumefaciens vir* genes, the *H. pylori cag* pathogenicity island, and *L. pneumophila dot/icm* gene regions. Colored arrows indicate recognized homologs in each T4SS. Genes encoding components of the presumed ‘core complex’ in each system are indicated by red text. (B) Protein sequence alignment of the VirD4 coupling proteins *H. pylori* Cag5 and *L. pneumophila* DotL. Protein sequences were aligned using BLAST align, and aligned sequences were visualized using Jalview. Numbers indicate the position of each residue in the pair-wise sequence alignment. Shading indicates level of sequence conservation.

## References

1. P. J. Christie, K. Atmakuri, V. Krishnamoorthy, S. Jakubowski, E. Cascales, Biogenesis, architecture, and function of bacterial type IV secretion systems. Annu Rev Microbiol 59, 10.1146/annurev.micro.1158.030603.123630 (2005).

2. E. Cascales, P. J. Christie, The versatile bacterial type IV secretion systems. Nature reviews. Microbiology 1, 137–149 (2003).

3. C. E. Alvarez-Martinez, P. J. Christie, Biological diversity of prokaryotic type IV secretion systems. Microbiol Mol Biol Rev 73, 775–808 (2009).

4. V. Chandran Darbari, G. Waksman, Structural biology of bacterial type IV secretion systems. Annu Rev Biochem 84, 603–629 (2015).

5. S. Odenbreit et al., Translocation of *Helicobacter pylori* CagA into gastric epithelial cells by type IV secretion. Science 287, 1497–1500 (2000).

6. W. Fischer, Assembly and molecular mode of action of the *Helicobacter pylori* Cag type IV secretion apparatus. FEBS J 278, 1203–1212 (2011).

7. D. Hofreuter, S. Odenbreit, R. Haas, Natural transformation competence in *Helicobacter pylori* is mediated by the basic components of a type IV secretion system. Mol Microbiol 41, 379–391 (2001).

8. E. Fernandez-Gonzalez, S. Backert, DNA transfer in the gastric pathogen *Helicobacter pylori*. J Gastroenterol 49, 594–604 (2014).

9. S. Backert, N. Tegtmeyer, W. Fischer, Composition, structure and function of the *Helicobacter pylori cag* pathogenicity island encoded type IV secretion system. Future Microbiol 10, 955–965 (2015).

10. M. G. Varga et al., Pathogenic *Helicobacter pylori* strains translocate DNA and activate TLR9 via the cancer-associated *cag* type IV secretion system. Oncogene, (2016).

11. S. Zimmermann et al., ALPK1- and TIFA-dependent innate immune response triggered by the *Helicobacter pylori* type IV secretion system. Cell Rep 20, 2384–2395 (2017).

12. T. L. Cover, M. J. Blaser, *Helicobacter pylori* in health and disease. Gastroenterology 136, 1863–1873 (2009).

13. M. Amieva, R. M. Peek, Pathobiology of *Helicobacter pylori*-induced gastric cancer. Gastroenterology 150, 64–78 (2016).

14. E. Cascales, P. J. Christie, Definition of a bacterial type IV secretion pathway for a DNA substrate. Science 304, 1170–1173 (2004).

15. H. H. Low et al., Structure of a type IV secretion system. Nature 508, 550–553 (2014).

16. R. Fronzes et al., Structure of a type IV secretion system core complex. Science 323, 266– 268 (2009).

17. J. E. Gordon et al., Use of chimeric type IV secretion systems to define contributions of outer membrane subassemblies for contact-dependent translocation. Mol Microbiol 105, 273–293 (2017).

18. T. Kubori et al., Native structure of a type IV secretion system core complex essential for *Legionella* pathogenesis. Proc Natl Acad Sci U S A 111, 11804–11809 (2014).

19. D. Ghosal, Y. W. Chang, K. C. Jeong, J. P. Vogel, G. J. Jensen, *In situ* structure of the *Legionella* Dot/Icm type IV secretion system by electron cryotomography. EMBO Rep 18, 726–732 (2017).

20. A. E. Frick-Cheng et al., Molecular and structural analysis of the *Helicobacter pylori cag* type IV secretion system core complex. mBio 7, (2016).

21. M. Rohde, J. Puls, R. Buhrdorf, W. Fischer, R. Haas, A novel sheathed surface organelle of the *Helicobacter pylori cag* type IV secretion system. Mol Microbiol 49, 219–234 (2003).

22. C. L. Shaffer et al., *Helicobacter pylori* exploits a unique repertoire of type IV secretion system components for pilus assembly at the bacteria-host cell interface. PLoS Pathog 7, e1002237 (2011).

23. T. Kwok et al., *Helicobacter* exploits integrin for type IV secretion and kinase activation. Nature 449, 862–866 (2007).

24. R. M. Barrozo et al., Functional plasticity in the type IV secretion system of *Helicobacter pylori*. PLoS Pathog 9, e1003189 (2013).

25. E. M. Johnson, J. A. Gaddy, B. J. Voss, E. E. Hennig, T. L. Cover, Genes required for assembly of pili associated with the *Helicobacter pylori cag* type IV secretion system. Infect Immun 82, 3457–3470 (2014).

26. L. F. Jimenez-Soto et al., *Helicobacter pylori* type IV secretion apparatus exploits beta1 integrin in a novel RGD-independent manner. PLoS Pathog 5, e1000684 (2009).

27. C. M. Oikonomou, G. J. Jensen, A new view into prokaryotic cell biology from electron cryotomography. Nat Rev Microbiol 14, 205–220 (2016).

28. K. Wallden, A. Rivera-Calzada, G. Waksman, Type IV secretion systems: versatility and diversity in function. Cell Microbiol 12, 1203–1212 (2010).

29. J. Tanaka, T. Suzuki, H. Mimuro, C. Sasakawa, Structural definition on the surface of *Helicobacter pylori* type IV secretion apparatus. Cell Microbiol 5, 395–404 (2003).

30. M. J. Kwak et al., Architecture of the type IV coupling protein complex of *Legionella pneumophila*. Nat Microbiol 2, 17114 (2017).

31. A. Redzej et al., Structure of a VirD4 coupling protein bound to a VirB type IV secretion machinery. EMBO J, (2017).

32. T. R. Costa et al., Structure of the bacterial sex F pilus reveals an assembly of a stoichiometric protein-phospholipid complex. Cell 166, 1436–1444 e1410 (2016).

33. S. Kutter et al., Protein subassemblies of the *Helicobacter pylori* Cag type IV secretion system revealed by localization and interaction studies. J Bacteriol 190, 2161–2171 (2008).

34. J. Andrzejewska et al., Characterization of the pilin ortholog of the *Helicobacter pylori* type IV *cag* pathogenicity apparatus, a surface-associated protein expressed during infection. J Bacteriol 188, 5865–5877 (2006).

35. S. Barden et al., A helical RGD motif promoting cell adhesion: crystal structures of the *Helicobacter pylori* type IV secretion system pilus protein CagL. Structure 21, 1931–1941 (2013).

36. E. M. Lai, C. I. Kado, Processed VirB2 is the major subunit of the promiscuous pilus of *Agrobacterium tumefaciens*. J Bacteriol 180, 2711–2717 (1998).

37. H. Schmidt-Eisenlohr et al., Vir proteins stabilize VirB5 and mediate its association with the T pilus of *Agrobacterium tumefaciens*. J Bacteriol 181, 7485–7492 (1999).

38. K. T. Pham et al., CagI is an essential component of the *Helicobacter pylori* Cag type IV secretion system and forms a complex with CagL. PLoS One 7, e35341 (2012).

39. T. Hayashi et al., Tertiary structure-function analysis reveals the pathogenic signaling potentiation mechanism of *Helicobacter pylori* oncogenic effector CagA. Cell Host Microbe 12, 20–33 (2012).

40. A. C. Shu et al., Evidence of DNA transfer through F-pilus channels during *Escherichia coli* conjugation. Langmuir 24, 6796–6802 (2008).

41. T. F. Anderson, Recombination and segregation in *Escherichia coli*. Cold Spring Harb Symp Quant Biol 23, 47–58 (1958).

42. T. F. Anderson, E. L. Wollman, F. Jacob, [Processes of conjugation and recombination in *Escherichia coli*. III. Morphological aspects in electron microscopy]. Ann Inst Pasteur (Paris) 93, 450–455 (1957).

43. W. F. Tivol, A. Briegel, G. J. Jensen, An improved cryogen for plunge freezing. Microsc Microanal 14, 375–379 (2008).

44. S. Q. Zheng et al., UCSF tomography: an integrated software suite for real-time electron microscopic tomographic data collection, alignment, and reconstruction. J Struct Biol 157, 138–147 (2007).

45. J. R. Kremer, D. N. Mastronarde, J. R. McIntosh, Computer visualization of three-dimensional image data using IMOD. J Struct Biol 116, 71–76 (1996).

46. J. I. Agulleiro, J. J. Fernandez, Fast tomographic reconstruction on multicore computers. Bioinformatics 27, 582–583 (2011).

47. D. Nicastro et al., The molecular architecture of axonemes revealed by cryoelectron tomography. Science 313, 944–948 (2006).

